## SupplementaryMaterials for "Anti-inflammatory effects of electrostimulation": Supplementary Materials.pdf

### Differential Anti-Inflammatory Effects of Electrostimulation in a Standardized Setting

##### Sampling Time points

Three time points were tested on almost all conditions (active and controls): baseline, 1h (early genes, as already performed in animal models (Nardini et al. 2016)). Regarding the final time point literature produces inconclusive consensus, owing to the extreme variability (and diversity from our) models:

- skin models are likely too immature to represent univoque model and coherent responses in this area. Research goes from the use of different types of electrostimulation on a variety of cell cultures or skin models to enhance wound healing assessing VEGF enhanced physiologic activation at 72h (Zhao et al. 2004). TGFB-pathway activation within 12h (Jennings, Chen, e Feldman 2008) and so forth, summarized along with additional punctual experiments in a review from 2014 (Yu, Hu, e Peng 2014).
- Most of human studies refer to the vagus nerve anti-inflammatory activity, not modeled in our system. One for all, reduction of TNF-alpha in Rheumatoid arthritis patients (chronic inflammation) was already significant after 4h by VNS (Koopman et al. 2016). Other works mention the nociceptive effects of electrical stimulation to last over one day (Goldman et al. 2010)

Personal communication from experts (Mike Cummings, MD and Luis Ulloa, Phd) indicated 48h as the time beyond which electrostimulation is not able to suppress TNF-alpha inflammatory activity (i.e.. after 48h the anti-inflammatory action of electrostimulation seems to be exhausted. For this reason the 3rd time point is chosen at 48h.

##### Electrical stimulation

Literature and personal communications (Mike Cummings MD, Vittoria Lauro MD, Luigi Manni PhD neurophysiologist) were used to reach consensus on the most appropriate experimental setting.

Stimulation can be released in the skin by: i) TENS patches (whose effects are comparable to needle stimulation (Ulett, Han, e Han 1998), limited however in our experimental setting by potential issue in adherence to the skin surface); ii) single point stimulation (HANS stimulation [xxxx], rejected for its limited diffusion in clinical practice); and iii) two needles stimulation by TENS machine, the latter was preferred as the most common therapeutic electro stimulatory approach. Stimulation is released by electrostimulator QiuTian Model SDZ II, and Tewa steel needles KJ-ART 0.20x25mm, following discussion with expert (Vittoria Lauro, MD, personal communication).

Parameters for the stimulation were defined again on a mixture of literature for the rationale, and medical experience for the relevance to clinical applications and future translatability. We decided to stimulate both in direct current (DC) and in alternate current (AC).

DC: endogenous electric fields occur naturally in vivo during wound healing and these are estimated to be 100–150 mV/mm at skin wounds (Song et al. 2007). To recreate a similar situation on our samples we decided to apply a voltage of 1V (DC) through an amplifier (trova la marca) and two needles at a distance of 0,8 cm for 20 seconds. Then we decided to raise up the voltage value to 5V (since the sample would burn at a higher voltage) to see if higher voltages could have an enhanced effect on the samples, b oth in the inflamed and physiologic samples.

AC: since electrostimulation on humans in the acupuncture treatments is performed in AC, we decided to try the voltage of 5V (to have the comparison with the direct current) and to try two different frequencies: a low frequency (10 Hz) and a high frequency (100 Hz) of electroacupuncture since it has been proven that high and low frequencies selectively induce the release of enkephalins and dynorphins in both experimental animals and humans (Han e Terenius 1982; Ulett, Han, e Han 1998).

**Methods for metabolomics**

The R package limma was selected for the differential abundance analysis of the metabolites, as its use can be generalized to more than 2 groups (in our case we had 57 analyzed contrasts) and it is quite powerful even for small-sized samples (Jeanmougin et al. 2010).

**Enrichment**

The function fgsea was used to perform a heterogeneous gene set enrichment analysis on multiple omics (Korotkevich et al. 2016). Other options were taken into consideration (i.e. MultiGSEA) but fgsea was preferred as it allows to integrate the metabolites into a single enrichment analysis, while MultiGSEA only allows to perform the analysis in parallel on transcripts and metabolites.
The results obtained from the differential analysis of the transcripts and metabolites using limma, and enriched using fgsea are in Supplementary Table S5-S6. The results of the subtraction of the enrichment results of the transcripts from the heterogeneous enrichment results are shown in Table S7.

Korotkevich, Gennady, Vladimir Sukhov, Nikolay Budin, Boris Shpak, Maxim N Artyomov, e Alexey Sergushichev. 2016. «Fast gene set enrichment analysis». BioRxiv, 060012.

Nardini, Christine, Valentina Devescovi, Yuanhua Liu, Xiaoyuan Zhou, Youtao Lu, e Jennifer E Dent. 2016. «Systemic wound healing associated with local sub-cutaneous mechanical stimulation». Scientific reports 6 (1): 39043.

Song, Bing, Yu Gu, Jin Pu, Brian Reid, Zhiqiang Zhao, e Min Zhao. 2007. «Application of direct current electric fields to cells and tissues in vitro and modulation of wound electric field in vivo». Nature protocols 2 (6): 1479–89.

Ulett, George A, Songping Han, e Ji-sheng Han. 1998. «Electroacupuncture: mechanisms and clinical application». Biological psychiatry 44 (2): 129–38.

Yu, Chao, Zong-Qian Hu, e Rui-Yun Peng. 2014. «Effects and mechanisms of a microcurrent dressing on skin wound healing: a review». Military Medical Research 1:1–8.

Zhao, Min, Huai Bai, Entong Wang, John V Forrester, e Colin D McCaig. 2004. «Electrical stimulation directly induces pre-angiogenic responses in vascular endothelial cells by signaling through VEGF receptors». Journal of cell science 117 (3): 397–405.
